# Effects of floral symmetry and orientation on the consistency of pollinator’s entry angle

**DOI:** 10.1101/2022.06.09.495424

**Authors:** Nina Jirgal, Kazuharu Ohashi

## Abstract

Since the publication of Sprengel’s (1793) observations, it has been considered that flowers with zygomorphic (or bilaterally symmetrical) corollas evolved to restrict the movement of pollinators into the flower by limiting the pollinator’s direction of approach. However, little empirical support has been accumulated so far, except Culbert and Forrest (2016) who found that zygomorphy reduced variance in pollinator’s flower entry angle. Our aim was to build on this work and observe whether floral symmetry or orientation had an effect on pollinator entry angle in a laboratory experiment using bumble bees, *Bombus ignitus*. Using nine different combinations of artificial flowers created from three symmetry types (radial, bilateral and disymmetrical) and three orientation types (upward, horizontal and downward), we tested the effects of these two floral aspects on the consistency of bee’s entry angle. Our results show that horizontal orientation significantly reduced the variance in entry angle, while symmetry had little effect. We also found no significant interactions between angle and symmetry in their effect on entry angle. Thus, our results suggest that horizontal orientation forces the bees to orient themselves relative to gravity rather than the corolla and stabilizes their flower entry. This stabilizing effect may have been mistaken for the effect of zygomorphic corolla as it is presented horizontally in most species. Consequently, we suggest that the evolution of horizontal orientation preceded that of zygomorphy as indicated by some authors, and that the reason behind the evolution of zygomorphy should be revisited.

## Introduction

Zygomorphy, or bilateral symmetry, in angiosperm flowers is suggested to have evolved independently in multiple lineages from their ancestral radial form about 50MY after their initial emergence, which coincides with the emergence of specialised pollinators (Citerne et al. 2010; Hileman 2014). Currently, 130 origins of zygomorphy have been estimated, while only 69 reversions to actinomorphy, or radial symmetry, have occurred (Reyes et al. 2016). Its emergence has been recognized as a key innovation as it is seen to be homoplastic in extant angiosperms and is associated with species diversification (Woźniak and Sicard 2018). Indeed, radially symmetric lineages comprise of fewer species than their bilaterally symmetric sister lineages (Sargent 2004; Woźniak and Sicard 2018). Overall, zygomorphic flowers have a higher level of fitness compared to their actinomorphic counterparts (Gómez et al. 2006).

Although there are many proposed hypotheses for the evolution of zygomorphy, they can be divided into three major groups of from the perspective of benefits for the plant. The first one is that zygomorphy may increase flower (re)visitation by pollinators through making the flowers easier to perceive, learn or forage. Because zygomorphic flowers are morphologically more complex than non-zygomorphic ones, pollinators are likely to be given more visual information on which to base their specific recognition of these flowers (Neal et al. 1998). For example, zygomorphic flowers have a higher contour density, i.e., the dissected margin of flowers, compared to actinomorphic ones (Anderson 1977; Dafni and Kevan 1997). Lehrer et al. (1985) have shown that honeybees tend to scan the contour of flowers in a close range. This may suggest that zygomorphic flowers exploit scanning behaviour of certain pollinators and help them to reach the floral resources (Dafni and Kevan 1997). Alternatively, zygomorphic flowers often have elongated lower lips on which pollinators could land easily (Sprengel 1793). The resultant decrease in landing time may lead to increased return visits by experienced foragers (Neal et al. 1998).

The second group of hypotheses is that the complexity of the corolla restricts the type of pollinators that could exploit the floral resources. Only a small subset of animals are able to reach the nectar in zygomorphic flowers due to their complexity (Neal et al. 1998; Zhao et al. 2016; Krishna and Keasar 2018). This would be advantageous if the complexity acts as a morphological filter that allows only effective pollinators to extract nectar from the flower and contribute to its pollination (Krishna and Keasar 2018). In addition, the filtering of pollinators according to their ability to handle flowers is suggested to increase pollinator fidelity, as specialised pollinators mainly exploit fewer complex flower species rather than visiting many flower species with simple morphologies (Rogriguez-Girones and Santamaría 2016; Krishna and Kaesar 2018). This fidelity of pollinators would benefit flowers in terms of lowering heterospecific pollen transfer (Krishna and Kaesar 2018).

The third hypothesis suggests that a zygomorphic corolla restricts the movement of the insect (Wang et al. 2014; Ushimaru and Hyodo 2005; Neal et al. 1998), resulting in approaches that are more stable and predictable in direction (Fenster et al. 2009). Zygomorphy has been linked to a gene that supresses the growth of stamen (Rudall and Bateman 2004) and concentrates the reproductive parts of the flower in one location. Indeed, O’Meara et al (2016) have shown that the diversification of zygomorphy is contingent on the presence of a corolla and reduction of stamen. The concentration of stamens and stigmas to a narrower area, combined with the predictable movement of the pollinator, would allow these reproductive parts to make more consistent contact with the pollinator’s body. This consistency would increase the conspecific pollen transfer while decreasing the heterospecific pollen exchange with pollinator-sharing species (Muchhala and Thomson 2010; Culbert and Forrest 2016).

Here we focus on the third hypothesis, which has often been assumed in studies of floral evolution (e.g., Sargent 2004) but rarely tested empirically. The effect of corolla shape on pollinator entry angle has been examined by Culbert and Forrest (2016) in a laboratory experiment using artificial flowers and bumble bees (*Bombus impatiens*). Using circular (radially symmetric) and rectangular (disymmetric) flowers, they showed that the approach consistency of bees was higher on disymmetric than on radial flowers. However, disymmetric flowers are rare in nature and slightly different from zygomorphic (bilaterally symmetric) flowers. In addition, all the artificial flowers in their experiment were oriented horizontally. While zygomorphic flowers are usually seen oriented horizontally in nature, radial flowers are commonly oriented in an upward or downward manner. Therefore, their results may not fully reflect the effect of corolla shape on entry angle consistency in natural settings. Finally, Fenster et al. (2009) has suggested that the entry angle stabilisation observed in zygomorphic flowers may be provided by the horizontal orientation in which they are presented, rather than the corolla symmetry. Thus, it is possible that the effects of floral symmetry were intermingled with those of orientation in the experiments by Culbert and Forrest (2016).

In this study, we aimed to evaluate both the effect of floral symmetry and orientation on the consistency of pollinator’s entry angle. As an extension of the experiment conducted by Culbert and Forrest (2016), we have used artificial flowers with three symmetry types (actinomorphy, zygomorphy and disymmetric) and oriented them in three ways (upward, horizontal and downward). By testing nine possible combinations of symmetry and orientation, some of which rarely exist in nature, we tried to dissect and quantify the effects of two floral features separately.

## Materials and methods

### Artificial flowers

We used artificial flowers for the experiments, each consisting of a “corolla” cut from blue drawing paper (∼11.5 cm^2^), and a container for cotton, from which the sucrose solution (“nectar”) can be collected by bees. Each corolla shape represents one of the three types of floral symmetry: actinomorphy (circular; 38 mm [diameter]), zygomorphy (triangular; 40 mm [base] ×57.5 mm [height]), and disymmetry (rectangular; 55 mm [length] × 21 mm [width]). We used artificial flowers with identical appearance for the training and test phases, the only difference being the configuration of the cotton container (see below).

A test flower had a 0.5-mL microcentrifuge tube as a container for a small piece of cotton. This microcentrifuge tube was embedded into a larger (1.5-mL) microcentrifuge tube using white, odourless clay. The opening of the smaller tube was at the same level as that of the larger tube. The cotton was inserted into the smaller tube, and the nectar was added on top of it so that bees could access it easily. The cotton was used as a fluid reservoir to prevent the nectar from leaking out of the microcentrifuge tube when the flower is not upwardly presented, while allowing bees to easily access it. On the other hand, a training flower used a 1.5-mL microcentrifuge tube as the container, into which a dental cotton roll was inserted to prevent the nectar from leaking out. We chose a larger nectar reservoir for the training flower, so that the nectar would not run out quickly and the bees would not lose their motivation. We also used six plastic petri dishes as supplementary feeders during the training phase (two petri dishes per flower shape). Each petri dish had a hole cut into the middle and a dental cotton was inserted. It absorbed the nectar in the petri dishes and allowed bees to access the nectar.

Twelve flowers were positioned on a grid in a three-by-four pattern, with a 10-cm interval between the centres of the adjacent flowers. Each was placed in either of the three positions: floor (grid lying flat on the floor of the cage), wall (grid leaning against the back side of the cage at 90° to the ground), and ceiling (grid hung upside down from the ceiling with metal hooks). Hereafter, we refer to these positions as “upward”, “horizontal”, and “downward” respectively, to represent how each position determines the flower orientation. Within each grid, we aligned the corollas so that their symmetry axes all point in the same direction to minimize the possible effects of variable corolla alignment. For upward and downward presentations, both the longer symmetry axes of disymmetric flowers and the symmetry axes of zygomorphic flowers were aligned parallel to the line connecting the nest and the cage. Moreover, the sharpest vertices of zygomorphic flowers were directed towards the wall opposite to where bees enter the cage. For horizontal presentations, the zygomorphic and disymmetric flowers were aligned so that an approaching bee could view them as upright triangles or upright rectangles, respectively (Fig. 1 d,e).

**Fig 1.**
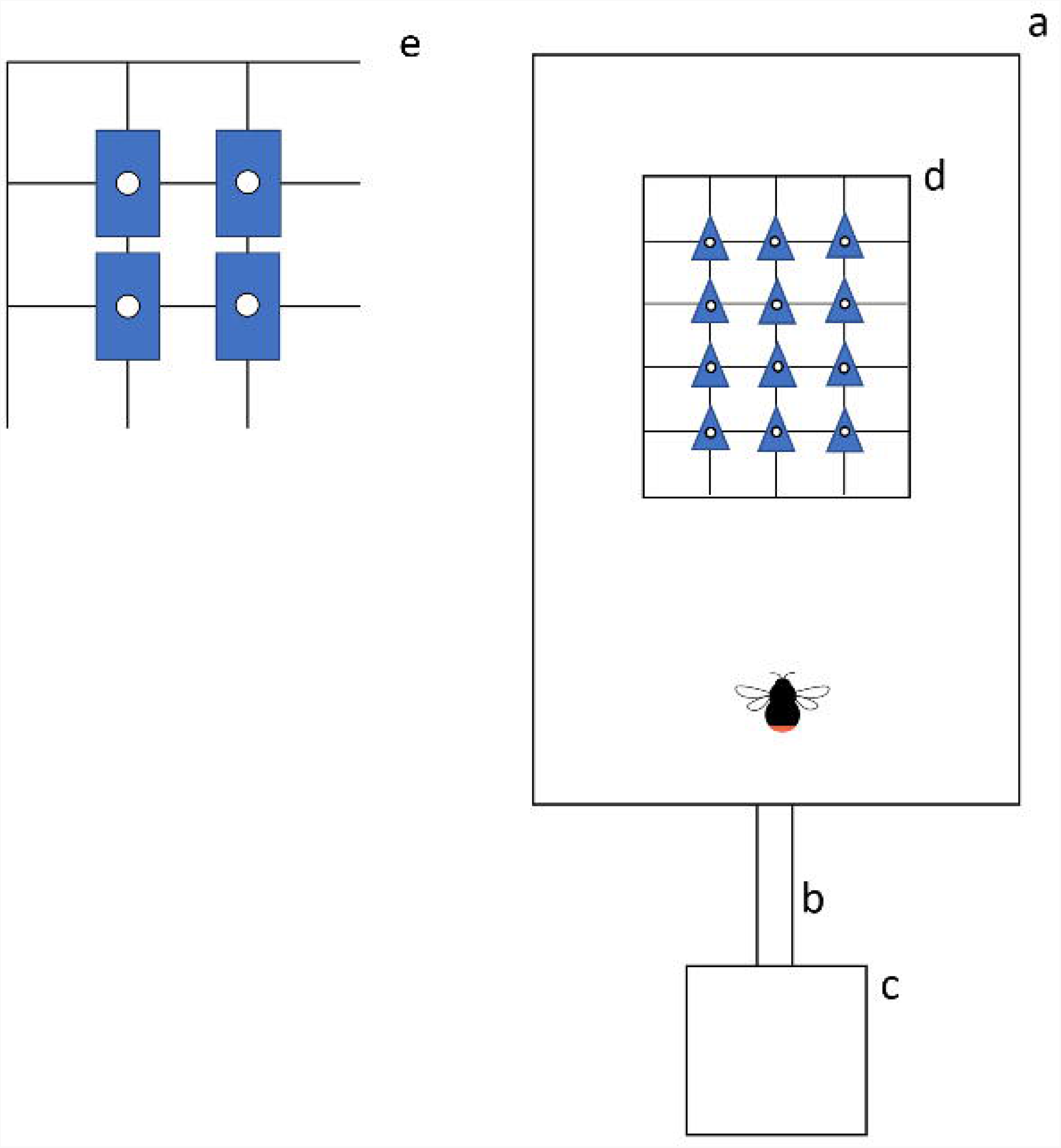
An aerial view of the experimental set up when flowers were oriented upwardly. Aa: flight cage; b: tunnel for bees to pass through with gated entrance; c: colony; d,e: grid of artificial flowers. The alignments is shown for zygomorphic (d) and dissymmetric (e) flowers, respectively.

#### Experimental procedures

We used workers from two commercial colonies of bumble bees, *Bombus ignitus* Smith, provided by Agrisect, Ibaraki, Japan. Colonies were maintained in nest boxes. The nest box (one at a time) was connected to a flight cage measuring 100 × 70 × 70 (H) cm through a transparent box equipped with gates, which allowed for the controlled entry and exit of individual bees into and out of the cage (Fig. 1). Pollen was supplied *ad lib* every day, directly into the colony. We used 17 and 18 workers from each colony, respectively.

During the training, which was performed before and between the test trials, we allowed the bees to forage freely in the cage by leaving the entrance open. Each training consisted of two phases: initial training phase and advanced training phase. During the initial training phase, a single training grid was placed on the floor of the cage (upward). In the grid, there were four flowers of each symmetry type arranged so that flowers of the same type were not next to each other. The initial phase was used to encourage bees to learn the association between the appearance of the corolla and nectar. Once bees started regular foraging on the training flowers, we proceeded to the advanced training phase, where two training grids were placed in the cage horizontally and downwardly, respectively. We also added six petri dishes (two per corolla shape), which were haphazardly placed on the floor as supplementary feeders. The training flowers and petri dishes were filled with 20% (w/w) sucrose solution and were replenished appropriately. Once consistent foraging began, we uniquely marked reliable foragers on their thorax with numbered, coloured tags.

Test trials were conducted using a single test grid placed in the cage. For each trial, we randomly selected one of the three symmetry types and arranged 12 of them in the grid, and then selected one of the three orientation types. In other words, one of the nine combination of symmetry and orientation was haphazardly chosen for each trial. As a reward, 10-μL of 30% sucrose solution was used (the concentration was increased to boost the bees’ motivation for foraging). A marked bee was selected haphazardly to carry out the trial. Bee foraging was filmed with a video camera (GZ-MG575-S, JVC Kenwood, Yokohama, Japan) that was placed at 90° to the flowers. When the bee failed to land on a flower for longer than five minutes or attempted to return to the nest through the gated entrance, we considered the foraging trip to be over; these bees were allowed to return to the nest on its own, or manually taken from the flight cage to the nest.

After the experiment, we went through the videotaped images and took a screenshot of each successful landing. A successful landing was defined as the bee’s posture being stopped on a flower and her proboscis extended. For each of them, the entry angle was measured using ImageJ (National Institute of Health, Version 1.52q, 2019). We measured the angle between two lines extending from the centre of the flower, i.e., one which goes vertically down the middle of the flower and another goes down the midline of the bee’s body. We defined the vertical line as a zero degree and measured the counter-clockwise angle of the midline of the bee’s body on a 0°–360° scales as the directionality of the bee’s entry (Fig. 2).

**Fig 2.**
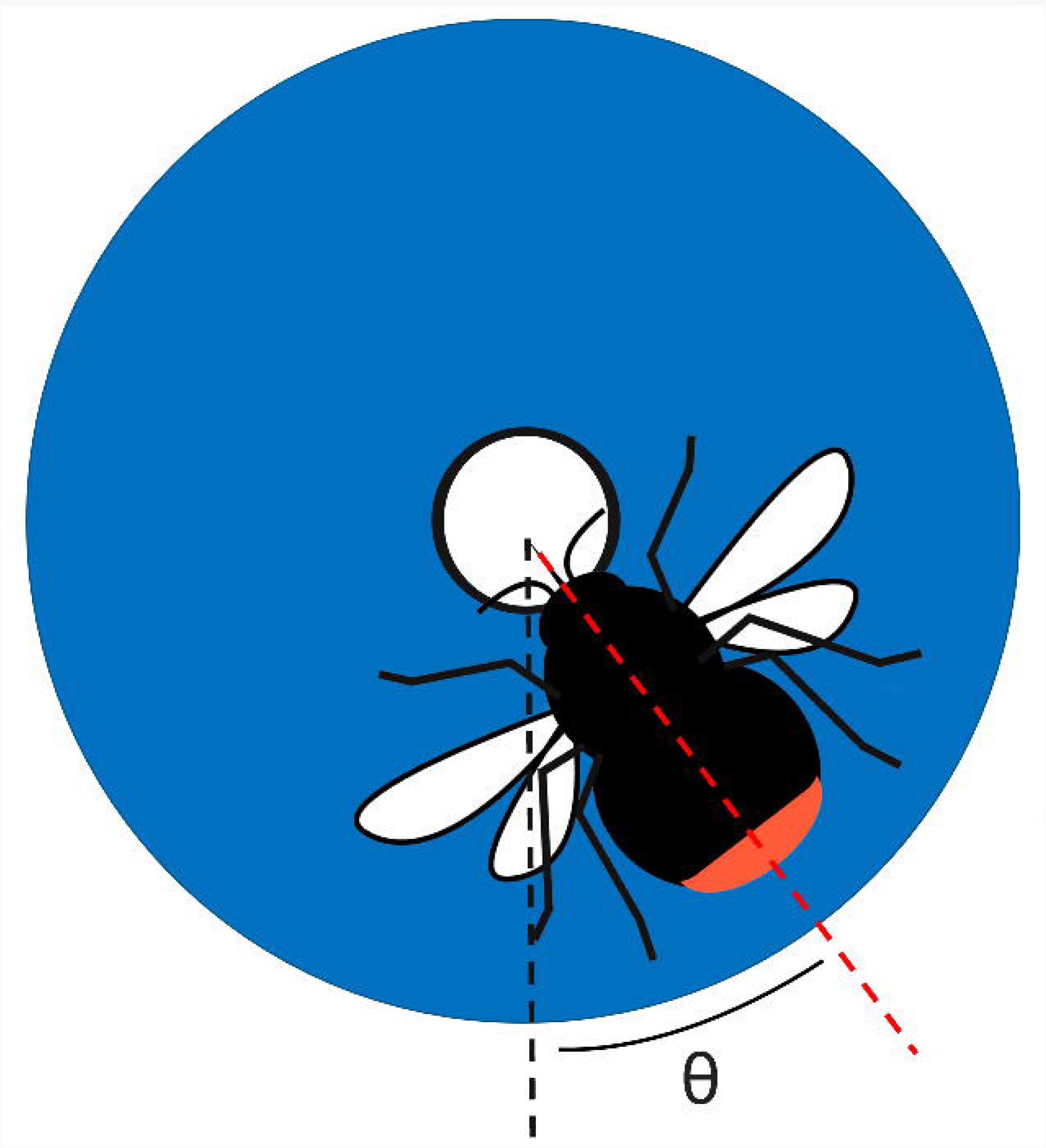
Schematic representation of the method used for measuring the entry angle of a bee. Two dotted lines were drawn from the centre of the artificial flower, one going down the middles of the flower (black) and another down the midline of the bee. We defined the black line as a zero degree and measured the counter-clockwise angle between these lines on a 0-360°scale as the bee’s entry angle.

We also measured the time it took for the bees to land on each type of flower (hereafter, “landing time”) to check if floral symmetry or orientation affects the ease of landing on flowers for bees. This was conducted by placing a video camera at about 90°, 50-80cm away from the flight cage, and recorded the sideview of the grid. We randomly selected three landings per trial and counted the number of frames it took for the bee to land on the flower, using Windows Media Player (30 frames per second). Because a bee usually initiates a hovering phase, approximately 8 mm away from the flower, before moving forwards to land (Reber et al., 2016), we started counting the frames when the bee first hovered in front of the target flower. The bee was considered to be hovering when it stayed in the same position for two or more frames. The counting was continued until the bee touched the flower with either of its legs and the legs remained on the flower during the successive two frames.

### Statistical analysis

We first converted the data of entry angle from degrees to radians. We then calculated the circular variance (mean resultant length, MRL) for each trial, as the ratio of the observed length of the resultant vector to the maximum possible length of resultant vector for the same size of sample. The maximum possible length of resultant vector is obtained when all the entries were in the same direction. For actual computation of MRL, we used the *rho*.*circular* function in the R package “circular” (Pewsey et al., 2013). The MRL will take a value from one to zero, which is difficult to interpret in terms of actual angles. Therefore, we converted the MRL into circular standard deviation (circular SD) using the formula:

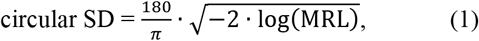

where π is the circular constant. The circular SD, like the usual standard deviation (SD), represents how much any one arbitrary data point deviates from the central value (mean entry angle) in unit of degrees (Pewsey et al., 2013).

We then fitted a linear-mixed model (LMM) to the data to determine whether and how floral symmetry and orientation affected the variance of bee’s entry angle. The circular SD was used as the response variable. We considered floral symmetry and orientation as fixed effects, bee individual as a random effect, together with an interaction term between symmetry and orientation. A type II Wald Chi-square test was performed to determine the significance of the fixed effects and the interaction.

Because we measured the landing time as the number of frames in video image, we fitted a generalised linear-mixed model (GLMM) to this data, using a logarithmic link function and a Poisson error distribution. We calculated the average number of frames elapsed for a landing in each trial, and then used it as a response variable. Floral symmetry and orientations were considered fixed effects, bee individual as a random effect, together with the interaction term between symmetry and orientation. A type II Wald Chi-square test was performed to determine the significance of the fixed effects and interaction.

## Results

We found that the circular SD was significantly affected by orientation (Fig. 3, χ^2^ = 235.09, P <0.0001, type-II Wald chi-square test), while it was hardly affected by symmetry or the interaction between symmetry and orientation (symmetry: χ^2^ = 4.13, P = 0.17; symmetry x orientation: χ^2^ = 4.21, P = 0.38, type-II Wald Chi-square test). The post-hoc test suggests that bees were significantly more consistent in their entry angle on horizontally presented flowers than on upwardly or downwardly presented ones (Fig. 3).

**Fig 3.**
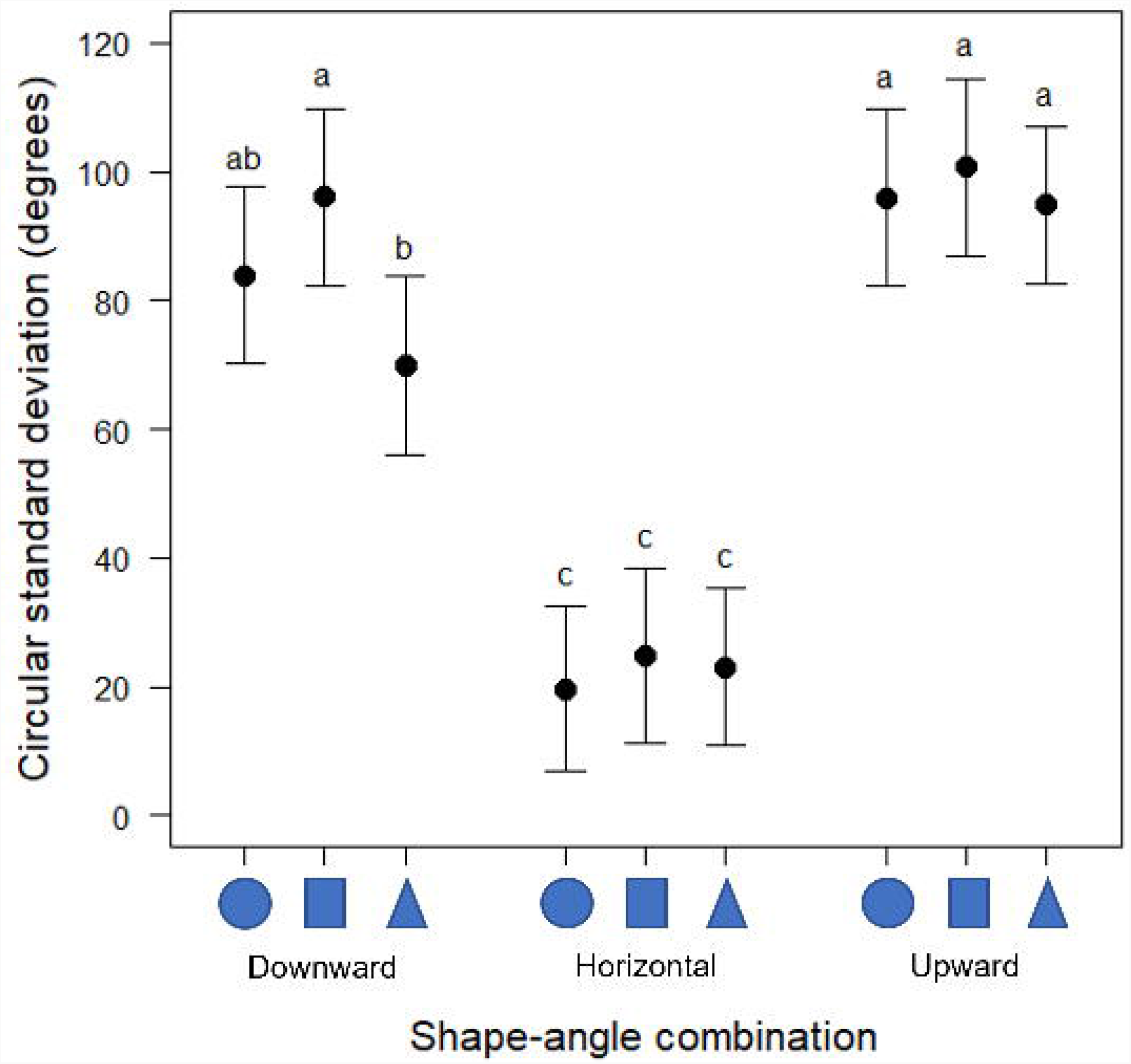
The circular SD of pollinator entry angle in nine shape-angle combinations. The shapes present on the X axis represent radial symmetry, bilateral symmetry, and zygomorphy, respectively. The data is divided into three groups; (a), (b), (c). Error bars indicate the standard error of the circular SD for each combination. Means with shared letters indicate that there is no significant difference at a 0.05 alpha level. Significance levels are adjusted by Tukey correction. The data indicates that horizontal orientation results in a lower circular SD, indication less variance in entry angle compared to other orientations. Upward and downward facing artificial flowers have a similar circular SD, however radial-downward and bilateral-downward showed slightly lower circular SD compared to other combinations of upward or downward flowers.

On the other hand, the landing time was significantly affected by orientation and the interaction between symmetry and orientation (Fig. 4, orientation: χ^2^ = 55.94, P <0.0001, symmetry x orientation: χ^2^ = 13.11, P = 0.011, type-II Wald Chi-square test), while it was hardly affected by symmetry (χ^2^ = 1.51, P = 0.47, type-II Wald Chi-square test). The post-hoc test suggests that bees took significantly longer landing on downwardly presented flowers than on upwardly or horizontally presented ones (Fig. 4).

**Fig 4.**
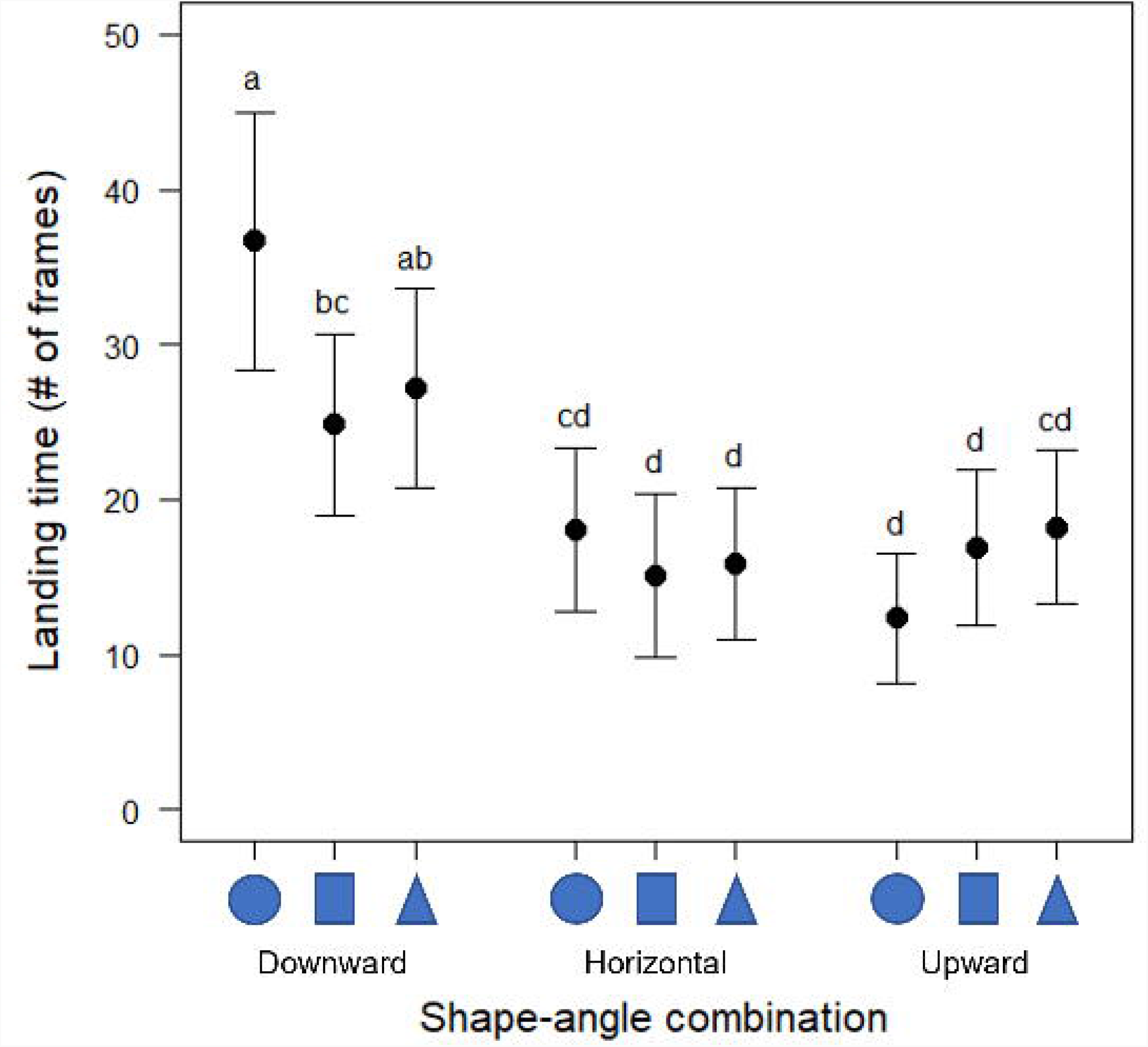
The average landing time (in number of frames) in nine shape-angle combinations. The shapes present on the X axis represent radial symmetry, bilateral symmetry, and zygomorphy, respectively. The data is divided into four groups; (a), (b), (c), (d). Error bars indicate the standard error of the circular SD for each combination. Means with shared letters indicate that there is no significant difference at a 0.05 alpha level. Significance levels are adjusted by Tukey correction. The data indicates that horizontal and upward facing flowers have lower mean landing times, while downward facing flowers had the longer landing times.

## Discussion

Bees were observed to enter horizontal flowers at a more consistent angle compared to the other two floral orientations (Fig. 3). In contrast, floral symmetry had little effect on the consistency of bee’s entry angle. The interaction between floral symmetry and orientation was not significant, either. We also found no significant effect of floral symmetry on the landing time of bees (Fig. 4). In addition, bees landed almost as quickly on both the horizontal and upwards flowers, while they took significantly longer to land on the downward flowers.

Since the publication of Sprengel (1793), it has often been assumed that zygomorphic corollas restrict the movement of a pollinator into the flower (e.g., Armbruster and Muchhala 2020). Based on this assumption, the pollen position hypothesis states that zygomorphy restricts the directionality of approach and movement of pollinators within and between flowers (Leppik 1972; Ostler and Harper 1978; Cronk and Moller 1997). However, our data shows that zygomorphy and disymmetry have little to no effect on the consistency of the bee’s entry angle (Fig. 3). Like the radially symmetrical flowers, these flowers allowed the bees to approach from various directions when orientated upwardly or downwardly, resulting in increased variability of their entry angle. Even when presented horizontally, zygomorphy and disymmetry did not increase the stability of the bee’s entry compared to actinomorphy (Fig. 3).

Our results are inconsistent with those of Culbert and Forrest (2016), who found that bumble bees entered disymmetric flowers at a higher consistency than they did for actinomorphic ones. Although we do not know the reason for this discrepancy, we could at least say that the stabilizing effect of disymmetric flowers found in Culbert and Forrest (2016) was much smaller than that of horizontal flowers in our study: the former found that standard deviations of entry angles were approximately 9° lower in disymmetric than in actinomorphic flowers; in contrast, the latter found that approximately 68° of difference in circular standard deviation of entry angles between horizontal and the other-oriented flowers (mean ± SE of circular SD at horizontal flowers = 22.5 ± 3.8°, df = 58; upward flowers = 97.2 ± 3.9°, df = 67; downward flowers = 83.3 ± 4.1°, df = 67, estimated from the fitted GLMM). In other words, the stabilizing effect of floral orientation was more than seven times stronger than that of floral symmetry demonstrated in Culbert and Forrest (2016).

The most probable reason why horizontal orientation had the strongest effect on the consistency of the bee’s entry angle (Fig. 3) is because gravity forced the bees to fly with the ventral side of their body facing the earth. Fenster et al. (2009) also observed that hovering hummingbirds showed more consistent approaches when flowers were presented horizontally than when the orientation was vertical (upward) or semi-pendant. These observations strongly suggest that the stabilisation effect does not come from visual guidance of the corolla shape or the existence of landing platforms on zygomorphic flowers, but the orientation of flowers relative to the gravity. It has long been known that in nature zygomorphic flowers are typically presented in a horizontal orientation (Ushimaru and Hyodo 2005). Therefore, much of our field impression that pollinators approach to zygomorphic flowers more consistently may have been heavily influenced or mislead by this strong correlation between zygomorphy and horizontality.

Our data supports the idea proposed by Fenster et al. (2009) that the stability of pollinator entry angle would be first conferred by the evolution of horizontal flowers that have been driven by abiotic stress, such as rainfall. They also pointed out the possibility that horizontal orientation set the stage for the evolution of symmetry in sexual organs, as a result of its stabilising effect on pollinator entry. It is likely that horizontal presentation of flowers promoted the evolution of symmetric sexual organs in flowers through the increased accuracy of pollen placement. Considering that corolla symmetry had little effect on the stability of entry angle (Fig. 3), however, it seems questionable whether this horizontality eventually lead to the evolution of zygomorphic corolla.

Our finding leaves open the question as to why zygomorphic flowers have evolved in the first place, especially in association with horizontal presentation. Given the current information, the most likely evolutionary advantage of zygomorphic corollas would be the restriction of pollinator type, to select for specialised and effective pollinators. This has been supported by empirical evidence (Lázaro and Totland 2014; Yoder et al. 2020). However, it remains unclear if the restriction of pollinator type works better when zygomorphic flowers are presented horizontally.

Alternatively, zygomorphic flowers might increase the ease of landing through visual guidance, leading to an increased attractiveness for pollinators. This was unsupported by our landing time data (Fig. 4), in which no significant association was detected between corolla symmetry and landing time. The idea that zygomorphic corolla increases the attractiveness was also unsupported, at least in bumble bees (Culbert and Forrest 2016). There is a possibility that the evolution of zygomorphy may be driven by natural selection acting on the lower lips, serving as a landing platform for pollinators, and the zygomorphic corolla could be a developmental by-product of this selection. Future studies should explore the precedence of horizontality in the evolution of zygomorphy, as well as the selective advantages of a zygomorphic corolla or its associated traits, such as well-developed lower lips. The use of 3D flowers will make it possible to test whether the existence of lower lips on zygomorphic flowers, which are more common in nature than the 2D flowers we used here, could change the results on the effects of the association between zygomorphy and horizontal orientation on the consistency of pollinator’s entry angle. Finally, we also urge future studies to quantify the effect of floral symmetry and orientation on pollen transfer to see how plausible the previous assumptions are in terms of actual pollination accuracy.

In sum, we first attempted to dissect the entangled effects of corolla symmetry and orientation on the consistency of pollinator’s entry angle. We presented compelling evidence that the visual symmetry of corolla has little effect on the consistency of entry angle. Rather, we found that horizontal presentation of flowers plays the largest role in stabilizing pollinator entry. These results may force a reconsideration of the common conception about the evolutionary significance of zygomorphic flowers. That is, zygomorphic corollas have often been assumed to stabilize pollinator’s entry to flowers. This could be a misconception caused by the fact that zygomorphic flowers are typically presented at horizontal orientation. That is, horizontal orientation, rather than corolla symmetry, may stabilize entry angle by forcing the pollinator to orient itself relative to gravity. We thus suggest that zygomoprhy in and of itself may not restrict the entry angle of pollinators but may instead allow for the evolution of corolla shape differences which further restrict the entry angle, thus leading to more precise pollen placement. In this respect, our data may support Fenster et al.’s (2009) proposal that horizontal orientation preceded the evolution of zygomorphy. Future studies are needed to determine how the stabilization effect of horizontal presentation affect pollination accuracy, as well as whether and how zygomorphic flowers have evolved and maintained in many angiosperm lineages.

## Declarations

### Funding

JSPS Grants-in-Aid for Scientific Research (KAKENHI no.19K06834)

### Availability of data

Data can be obtained from Nina Jirgal upon request.

## Acknowledgements

We would like to thank Kohei Terada and Yukie Sato for their insightful advice during the development of the experimental method; Ben Chapman, Rob Sansom, and William Sellers for their critical comments following assignments related to this experiment which were used in this study and Nathan Muchhala for his helpful comments and insights on the manuscript. This work was supported by a JSPS Grants-in-Aid for Scientific Research (KAKENHI no.19K06834) to K.O.

## Notes

### Competing Interest Statement

The authors have declared no competing interest.

